# “Isolation and characterization of a novel hormone receptor positive mammary adenocarcinoma MCa-P1362 with stromal drivers of tumor growth, metastasis, and drug resistance”

**DOI:** 10.1101/2023.06.02.543434

**Authors:** Samir Jana, Wende Li, Pin-Ji Lei, Zixiong Wang, Peigen Huang, Dennis Jones

## Abstract

Preclinical models that display spontaneous metastasis are necessary to improve therapeutic options for hormone receptor positive breast cancers. In this study, we conducted a detailed cellular and molecular characterization of MCa-P1362, a novel syngeneic Balb/c mouse model of metastatic breast cancer. MCa-P1362 cancer cells expressed estrogen receptors (ER), progesterone receptors (PR), and HER-2 receptors. MCa-P1362 cells proliferate in vitro and in vivo in response to estrogen, yet do not depend on steroid hormones for tumor progression. Further characterization of MCa-P1362 tumor explants shows that they contain a mixture of epithelial cancer cells and stromal cells. Based on transcriptomic and functional analyses of cancer and stromal cells, stem cells are present in both populations. Functional studies demonstrate that crosstalk between cancer and stromal cells promotes tumor growth, metastasis, and drug resistance. MCa-P1362 may serve as a useful preclinical model to investigate the cellular and molecular basis of hormone receptor positive tumor progression and therapeutic resistance.

## Introduction

Breast cancer accounts for 30% of newly diagnosed cancers and is the second leading cause of cancer-related death among women ^1, 2^. Most breast cancers are driven by hormone receptors estrogen receptor alpha (ERα) and/or progesterone receptor (PR) ^3^. More than 70% of diagnosed breast cancers are ER+ ^4^. Many ER+ tumors depend on estrogen for growth and endocrine therapies targeting estrogen production, degradation of ER or modulation of ER signaling are often the first line of treatment for ER+ breast cancer ^5^. Despite initial responsiveness to endocrine therapies ^6^, about half of advanced (local or regional relapse, distant metastasis) ER+ cancers exhibit intrinsic and/or acquired resistance to such therapy ^7^. For example, over 20% of ER+ patients treated with tamoxifen, a selective inhibitor of estrogen receptor signaling, will develop recurrence within 15 years of diagnosis ^8^. Patients with metastatic ER+ breast cancer who experience distant recurrences have a median survival of approximately four years ^9^. New therapies are urgently needed since most breast cancer-related deaths are caused by metastasis ^10^. Preclinical research to study ER+ tumors has been hindered by the lack of metastatic ER+ mouse breast cancer models, even though most breast cancers are ER+. Many genetically engineered mouse models are available that include constitutive or inducible transgenic approaches for modeling tumor progression and metastasis ^11^. However, most do not reliably develop metastases and thus limit our understanding of the key molecular targets for therapy after acquired resistance to targeted and systemic therapies.

The ER+ human T47D and MCF-7 cell lines are commonly used to interrogate the response to endocrine agents. Additionally, patient-derived xenograft (PDX) models have demonstrated efficacy in predicting patient response to anticancer therapies but have mainly allowed interrogation of ER-cancers since they have a higher establishment rate than ER+ cancers ^12^. Moreover, human breast cancer lines and PDX models depend on exogenous 17-β estradiol for growth and subsequent metastasis. Yet, most ER+ breast cancers arise in postmenopausal women with low estrogen levels ^13^, making induction of high estrogen levels less clinically relevant.

An additional limitation of xenograft models is their ability to assess tumor-stromal interactions since human stroma is lost after transplantation and replaced with mouse cells and protein not orthologous with human. Notably, mesenchymal cells, which are the predominant cell type in the breast tumor stroma ^14^, promote metastasis, immunosuppression, tumor progression, and therapeutic resistance. For example, cancer-associated fibroblasts (CAFs), a type of mesenchymal cell found in breast tumors, secrete cytokines and growth factors, as well as extracellular vesicles that reduce ER expression and tamoxifen sensitivity, thus contributing to endocrine therapy resistance ^15, 16^. CAFs have also been shown to promote the survival of breast cancer stem cells (CSCs) ^17^, which play a central role in therapeutic resistance. As an example, CSCs are significantly enriched in ER+ MCF-7 cells *in vitro* and in patient-derived xenografts treated with endocrine therapy ^18^. Furthermore, the *in vivo* study of human cancer cells requires mice that lack a functioning immune system, thus preventing our full understanding of the interaction between cancer cells, stromal cells, and immune cells. To test the development of novel therapies for ER+ breast cancers that develop resistance to current therapies and to understand how ER+ breast tumors evade anti-tumor immunity, syngeneic mouse ER+ breast models are needed, as xenograft models may not capture the complex intra-tumor heterogeneity that dictates immune evasion, tumor progression and metastasis.

In this study, a syngeneic murine breast cancer line with characteristics of luminal B (ER + , PR + and HER2 +) subtype was established and characterized *in vivo* and *in vitro*. MCa-P1362 cancer cells grow without estrogen supplementation and spontaneously metastasize to multiple organs in syngeneic Balb/c mice. This hormone receptor positive breast cancer model recapitulates mechanisms of endocrine therapy resistance conferred by the tumor stroma and by cancer stem cells (CSCs).

## Results

### Spontaneous breast adenocarcinoma with metastatic outgrowth

In our defined-flora animal facility ^19–21^, a 339-day-old female syngeneic Balb/c breeder developed a spontaneous tumor of the right chest. The body weight of the tumor-bearing host was 35.5 grams. Following euthanasia and necropsy of the mouse, a grossly visible subcutaneous mass of 1.7 x 1.4 x 1.3 cm localized in the second mammary fat pad (MFP) region was revealed (P1362). The tumor was well-circumscribed, firm, and multicolored – white, gray, and yellow on cut surface. This tumor also presented with central necrosis and hemorrhage, as well as local invasion into the surrounding normal tissues, but without distant lung or liver metastasis. This spontaneous adenocarcinoma, P1362, was then transplanted and serially passaged in the MFP of female Balb/c mice for 5 generations (F1-F5) (**Fig. 1A**). The P1362 Isografts grew in 95% (19/20) of transplanted hosts with an average latent period of 9.5 ± 7.7 days and growth time of 45.0 ± 18.4 days (**Table 1**). F1 tumors had a latent period of approximately 26 days and growth time of approximately 85 days (**Fig. 1B**). F2-F5 isografts grew faster with a latent period of approximately 6 days and growth time of approximately 37 days. Histological examination revealed a typical adenocarcinoma feature (type B) with a glandular growth pattern (**Fig. 1Ci**). All P1362 transplanted tumors were histologically similar to the original adenocarcinoma (**Fig. 1Dii-iv**).

**Figure 1.**
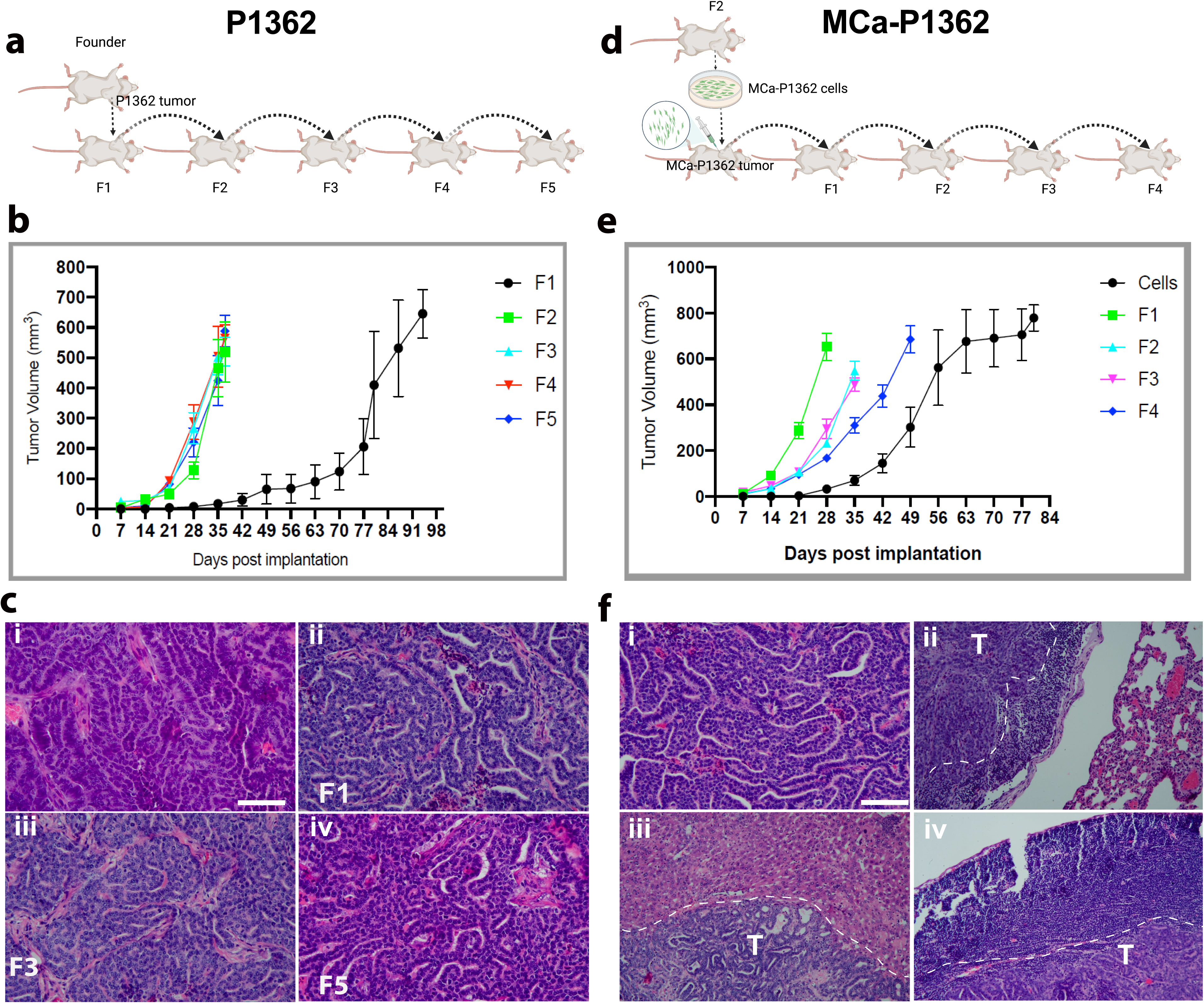
Isolation and propagation of a spontaneous breast tumor from a Balb/c mouse. **(A)** Schematic of in vivo passaging from original P1362 tumor. (**B**) Tumor growth kinetics of P1362 mammary tumors transplanted into mammary fat pads of Balb/c mice and serially passaged *in vivo* from F1 to F5. **(C)** Hematoxylin and eosin staining of tissue sections illustrating the histopathology of the original P1362 spontaneous mammary tumor in a Balb/c mouse, demonstrating typical adenocarcinoma histological features (type B) accompanied by a glandular growth pattern (**i**), as well as P1362 isografts (**ii**, F1; **iii**, F3; and **iii**, F5) in Balb/c mice that exhibit similar histological characteristics as the original P1362 adenocarcinoma. (**D**) Schematic of *in vivo* passaging from MCa-P1362 cells derived from P1362 F2 tumors. (**E**) Tumor growth kinetics of mammary tumors derived from MCa-P1362 cells injected into mammary fat pads of Balb/c mice. Tumors were then serially passaged *in vivo* from F1 to F4. **(F)** Images of hematoxylin and eosin-stained tissue depicting the histological features of the MCa-P1362 MFP isograft in Balb/c mice that resemble those of the original spontaneous tumor P1362 (**i**). MCa-P1362 isograft metastases to lung (**ii**), liver (**iii**), and lymph node (**iv**) in Balb/c mice (T: tumor; Scale bar = 100 µm. Data are presented as mean tumor volume; bars depict SEM. n=4-5 per group.

Next, fresh tumor tissues from P1362 F2 isografts were harvested and cultured, as previously described ^19–21^, to obtain an *in vitro* MCa-P1362 cell line and to establish an *in vivo* MCa-P1362 tumor line (**Fig. 1D**). 1x 10^6^ MCa-P1362 cells were then injected into the third MFP of female Balb/c mice to grow isografts (MCa-P1362 cells). Fragments of the isografts that developed from injected MCa-P1362 cells were then transplanted and serially passaged in female Balb/c mice for 4 generations (MCa-P1362 F1-F4). MCa-P1362 tumors had a 100% (18/18) tumor take rate and an average latent period of 8.4 ± 5.8 days and growth time of 39.6 ± 12.8 days (**Table 1**, **Fig. 1E**). Unlike the P1362 F1-F5 tumor bearing mice that did not develop spontaneous metastases, we found that 50% (9/18) of MCa-P1362 isografts (after surgical removal of the MFP primary tumors at 3 months ^20^) metastasized to local lymph nodes, lungs, and livers. Among these 9 cases, five cases involved lymph nodes with lung or liver metastases, and four cases involved pulmonary metastases only. All MCa-P1362 transplanted tumors closely resembled the original adenocarcinoma histopathological features (**Fig. 1Fi**). Their metastases were also confirmed histologically (**Fig. 1Fii-iv**).

### Hormone receptor status of MCa-P1362 tumor cells

By western blotting, we assessed protein expression levels of the hormone receptors ERα, PR, and the growth factor receptor HER-2 in MCa-P1362 tumor cells. MCa-P1362 cells were compared with 4T1, BT474, and MCF-7 breast cancer cells for the expression of these receptors. MCa-P1362 cells expressed ERα at levels comparable to ERα-positive MCF-7 human breast cancer cell line (**Fig. 2A**). Surprisingly, 4T1 cells showed low expression of ERα although it has been described as a triple negative model of breast cancer ^22^. HER-2 was also expressed in MCa-P1362 tumor cells, but at levels lower than the HER-2 enriched BT-474 human breast cancer cell line. Progesterone receptors were expressed in the cell lines probed except 4T1. By immunofluorescence, ERα, PR, and HER2 expression was also detected in cultured MCa-P1362 cells (**S.** **Fig. 1A**). We next assessed the functionality of ERα by measuring the growth of MCa-P1362 cells in the presence of 17-β estradiol, an ERα ligand. To this end, MCa-P1362 cells were treated with increasing concentrations of estradiol for up to 4 days. We found that estradiol stimulated MCa-P1362 cancer cell proliferation in a dose-dependent manner, although high concentrations (>10 nM) of estradiol exposure were toxic to MCa-P1362 cancer cells (**Fig. 2B**). To measure whether ERα signaling stimulates MCa-P1362 tumor growth, we evaluated ERα expression in MCa-P1362 tumors by immunofluorescence. We measured a heterogeneous pattern of expression, with 16% of cytokeratin-positive cancer cells in MCa-P1362 tumors expressing ERα 54 days after MCa-P1362 cell implantation (**S** **Fig. 1B**). In MCa-P1362 tumors, HER2-positive cancer cells were also less prevalent than in culture conditions (**S** **Fig. 1A**), while PR levels in MCa-P1362 tumors remained comparable to MCa-P1362 cells in culture **(S.** **Fig. 1B**). To determine the effects of estradiol on MCa-P1362 tumor development, we implanted slow-release estradiol pellets at the time of MCa-P1362 cell injection into mice. Estradiol supplementation significantly increased the growth of MCa-P1362 tumors compared to placebo pellets (**Fig. 2C**). Further, estradiol increased the incidence of lymph node metastasis from 25% to 75% (**Fig. 2D**). Tamoxifen, an estrogen receptor antagonist, is a mainstay of treatment for patients with ER+ breast cancer ^23^. We next determined if tamoxifen could slow the growth of MCa-P1362 cells. To this end, MCa-P1362 cells were exposed to increasing doses of 4-Hydroxytamoxifen (4-OHT) and cell viability was measured. MCF-7 cells were used as a positive comparator. Based on MTT viability assays, 4-OHT inhibited the growth of both MCa-P1362 and MCF-7 cells *in vitro* with an IC_50_ of 3.6 µM and 0.9 µM, respectively (**Fig. 2E**) at 24 hrs. Interestingly, MCa-P1362 cells were more resistant to the cytotoxic effects of 4-OHT compared to MCF-7 cells. 4-OHT, however, had a modest effect on the viability of 4T1 cells (**Fig. 2E**) at low concentrations (< 8 µM). We next established 4T1 and MCa-P1362 tumors and examined whether tamoxifen reduced their growth. Based on prior studies ^24^ of MCF-7 tumors, animals were treated daily with 100 µg tamoxifen. Compared to vehicle (corn oil)-treated animals, tamoxifen did not significantly affect the growth of primary 4T1 tumors (**Fig. 2F**). Tamoxifen slightly reduced MCa-P1362 primary tumor size (**Fig. 2G**), a trend that did not reach statistical significance. To evaluate the effect of endogenous estrogen and progesterone on MCa-P1362 tumor growth, MCa-P1362 cancer cells were implanted into ovariectomized syngeneic mice. Interestingly, the growth kinetics of MCa-P1362 tumors remained unaltered despite the ovariectomy (Fig. **Fig. 2H**). Together, these data demonstrate that MCa-P1362 cancer cells are responsive to, but not dependent on estrogen for growth.

**Figure 2.**
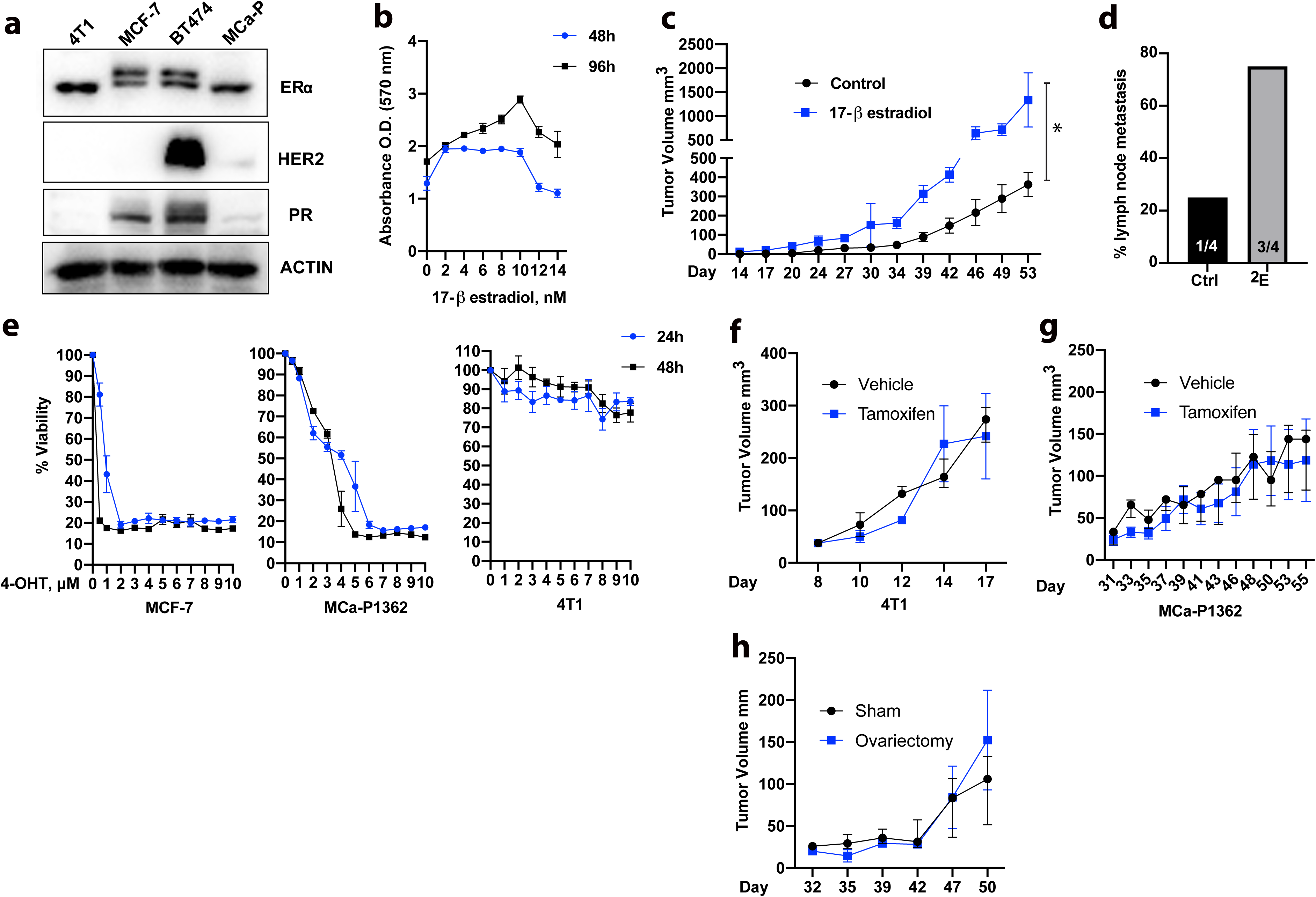
MCa-P1362 hormone receptor expression and response to estrogen therapy. **(A)** Detection of estrogen receptor (ERα), human epidermal growth factor receptor 2 (HER2), and progesterone receptor (PR) by western blot with lysates from 4T1, MCF-7, BT474, and MCa-P1362 (MCa-P) breast carcinoma cells. Actin was used as loading control. (**B**) MTT assay depicting proliferation of MCa-P1362 cells at 48 and 96 hours after exposure to indicated concentrations of 17-β estradiol. (**C**) Tumor growth curves of mice implanted with placebo pellet (control) and mice implanted with 17-β estradiol pellet (0.36mg) n=4 each group. Measurements were taken every 3-4 days once tumors became palpable. Each data point is the mean tumor volume. Data depicted as standard deviation. * *p* =0.02 by two-way ANOVA. (**D**) Incidence of lymph node metastases in after growth of MCa-P1362 tumors in the presence of placebo (Ctrl) or estradiol (E_2_)-pellets, as determined by cytokeratin staining of serial lymph node sections. (**E**) MCF-7, MCa-P1362, or 4T1 cells were treated with indicated concentrations of Hydroxytamoxifen (4-OHT) for 24 and 48 hours. Tumor growth curves of 4T1 (**F**) and MCaP-1362 (**G**) tumors (*n* = 4 for 4T1; n=5-6 for MCa-P1362) in response to control (corn oil) and tamoxifen (TAM) treatment. TAM treatment was started when the tumors reached approximately 30 mm^3^. (**H**) Effect of ovariectomy on MCaP-1362 tumor growth. n = 4/group.

### Heterogeneous primary culture of MCa-P1362 tumors

In many cancers, including breast carcinoma, cancer-associated fibroblasts (CAFs) make up most of the tumor stroma and their abundance is often associated with tamoxifen resistance and poor outcomes ^16, 25^. We performed immunofluorescence staining on MCa-P1362 tumors using antibodies against α-smooth muscle actin (SMA), a marker for CAFs^26^. The staining revealed the presence of CAFs in MCa-P1362 tumors (**Fig. 3A**), and their prevalence and distribution were similar to that of CAFs in human breast cancer (**Fig. 3B**). An enzymatic digestion was then used to dissociate tumor and stromal cells from MCa-P1362 tumors and establish primary cell cultures. Characterization by flow cytometry of cultured MCa-P1362 tumor explants revealed cells that were positive or negative for the epithelial marker epithelial cellular adhesion molecule (EpCAM) (**Fig. 3C**). EpCAM+ and EpCAM-cells were then sorted by FACS and plated. EpCAM-cells had spindle-shaped morphologies, typical for fibroblasts (**Fig. 3D**), while EpCAM+ cells demonstrated a cobblestone morphology (**Fig. 3E**). Immunofluorescence microscopy of sorted cells revealed that EpCAM+ cells stained positive for the cancer cell marker cytokeratin (**Fig. 3F**), but not the CAF marker α-SMA. Conversely, EpCAM-cells stained positive for α-SMA, but not cytokeratin **Fig. 3G**. As expected, EpCAM-cells exhibited slower growth than EpCAM+ cells over a 7-day period (**Fig. 3H**). In MCa tumors, cancer cells, but not CAFs were proliferative **(S** **Fig. 1C**).

**Figure 3.**
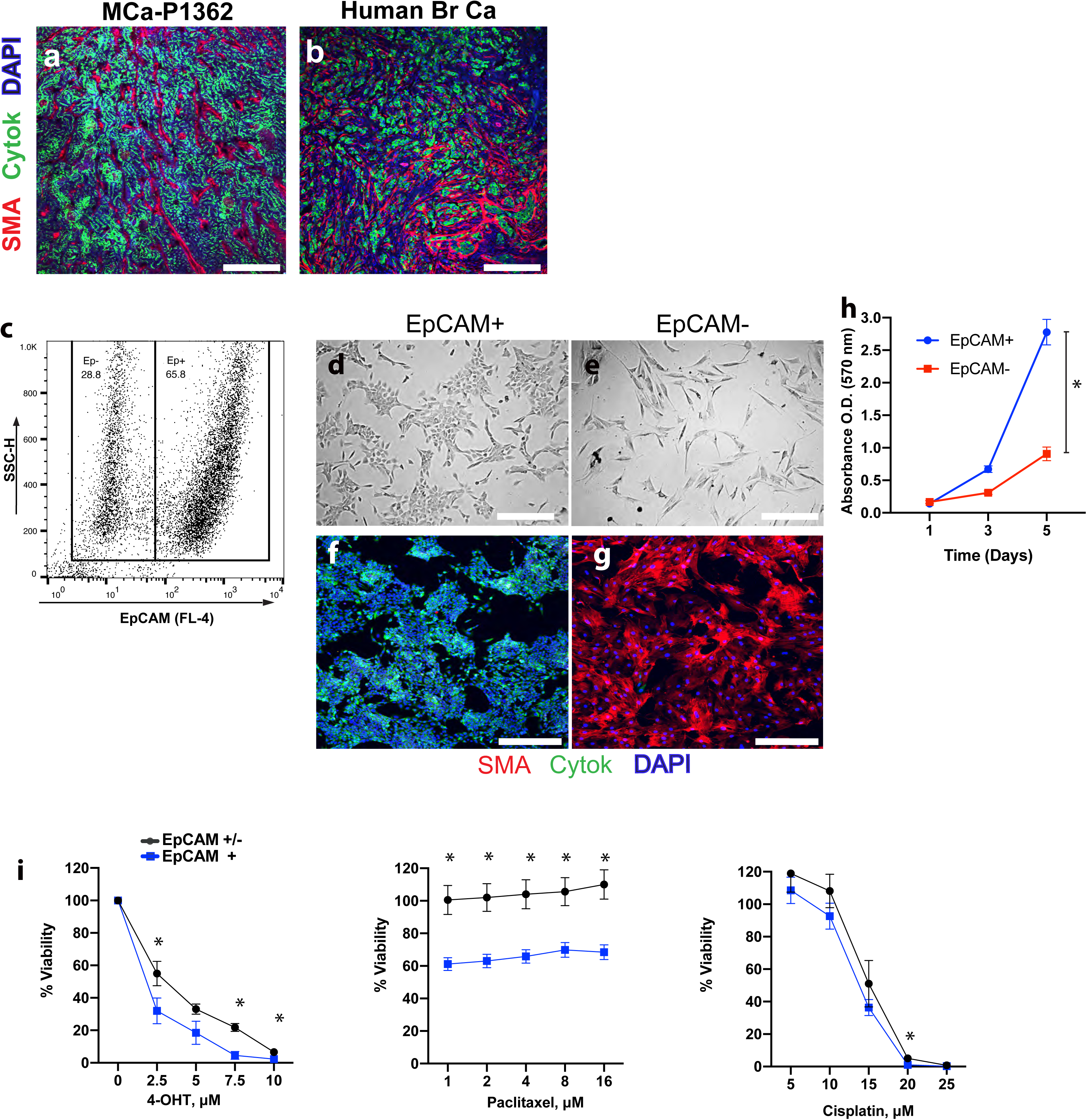
Characterization of distinct cellular populations from MCa-P1362 tumors. **(A-B)** Immunofluorescent staining of primary breast tumors. MCa-1362 tumor (A) and human tumor (B) were stained with antibodies detecting α-smooth muscle actin (SMA) (red)) and cancer cells (cytokeratin, green). Scale bars = 150 μm. (**C**) Flow cytometry dot plot represents the distribution of EpCAM- and EpCAM+ cells from MCa-P1362 culture. (**D, E**) Brightfield image of cells isolated from MCa-P1362 primary tumor and FACS-sorted based on EpCAM gating for flow cytometry. Scale bar = 300 μm, (**F,G**) Immunofluorescent staining of EpCAM+ and EpCAM-cells that were flow-sorted from MCa-P1362 culture. Cells were stained with antibodies against EpCAM (**F**) and alpha smooth muscle actin (SMA) (**G**). Scale bar = 300 μm. (H) EpCAM+ and EpCAM-cell proliferation evaluation over 5 days using MTT assay. *p<0.001 with two-way ANOVA. (I)Viability assay of sorted EpCAM+ cells or epithelial/mesenchymal (EpCAM+/-) co-culture with increasing doses of 4-OHT (left panel), paclitaxel (middle panel), and cisplatin (right panel). Measurements were taken 72 hours after cells were incubated with drugs. Data are displayed as % viability and normalized to untreated controls. 4-OHT * *p<*0.05. Paclitaxel * *p=* 0.0001. Cisplatin p=0.004. n=3. Significance was measured using two-tailed unpaired Student’s *t*-tests.

CAFs have been shown to confer resistance to breast cancer cells exposed to endocrine therapy or chemotherapy ^16, 17, 27^. To measure if CAFs from MCa-P1362 primary cultures are protective against these therapies, co-cultured EpCAM+ and EpCAM-cells or mono-cultures of sorted EpCAM+ cells were treated with 4-OHT, paclitaxel, or cisplatin for 72 hours (**Fig. 3I**). Co-cultures of EpCAM+ and EpCAM-cells showed nearly a 5-fold increase in viability after tamoxifen treatment (7.5 µM) compared to tamoxifen-treated Ep-CAM+ monocultures. Cancer cells were not responsive to paclitaxel in the presence of EpCAM-cells, whereas EpCAM+ monocultures displayed a greater than 40% decrease in viability after paclitaxel challenge. Although most MCa-P1362 cancer cells were killed with 20 µM cisplatin, EpCAM-cells provided moderate protection against its cytotoxic effects at all concentrations, with a significant effect measured at 20 µM ( 5% viability in co-culture compared to 1% viability in monoculture). (**Fig. 3I**). Collectively, these data suggest that EpCAM+ and EpCAM-cells represent cancer cells and CAFs, respectively, and that CAFs maintain their tumor-promoting properties ex vivo.

### Bulk RNA sequencing of MCa-P1362 tumors

To gain further insight into the MCa-P1362 cell line, we performed RNA sequencing (RNA-seq) of EpCAM+ and EpCAM-cells. MCa-P1362 cells were FACS-sorted into Ep-CAM+ or EpCAM populations and grown as monocultures before RNA isolation. Alternatively, MCa-P1362 cells were FACS-sorted into EpCAM+ or EpCAM populations and immediately frozen for RNA isolation (co-culture). Bulk RNA sequencing of the samples was then performed and subsequently analyzed for differential gene expression with the cutoff of ≥1.5-fold change and a false discovery rate ≤ 0.1. A total of 2648 differentially expressed genes (DEGs) were identified between the pre-sorted monocultured EpCAM+ and EpCAM-cells. A total of 2449 DEGs were identified between the EpCAM+ and Ep-CAM-cells that were cocultured (**Fig. 4A**). Among these differentially expressed genes, 1871 genes were overlapped, accounting for 58% of the total DEGs. An unsupervised clustering of the gene expression of these 3226 DEGs suggested that the monocultured pre-sorted cells maintained similar gene expression pattern compared to the cocultured cells (**Fig. 4B**). Gene ontology analysis of the DEGs between monoculture EpCAM+ and EpCAM-cells revealed expression of genes that were mainly involved in adhesion, migration, proliferation, extracellular matrix, and development (**Fig. 4C**). GSEA analysis demonstrated that genes related to epithelial development were upregulated in the Ep-CAM+ cells and genes related to extracellular matrix were enriched in EpCAM-cells (**Fig. 4D,E**). Using a pan-cytokeratin antibody, immunofluorescence revealed that EpCAM+ cells expressed cytokeratin, an epithelial marker (**Fig. 3F**). By bulk RNA-Seq, several cytokeratin transcripts were identified in the EpCAM+ cell transcriptome (**Fig. 4F**), including *Krt 7, 8, 18*, and *19*, associated with luminal epithelium and *Krt* 14 and 17, associated with basal epithelium. Similar to immunofluorescence staining (Fig. 3G), cytokeratin genes were not expressed in EpCAM-cells (**Fig. 4G**). However, fibroblast-associated genes including *S100A4* (fibroblast-specific protein 1), *Postn*, and *Acta2* were expressed in EpCAM-cells (**Fig. 4G**). Finally, quantitative PCR analysis of magnetically separated EpCAM+ and EpCAM-cells validated the expression of epithelial genes in EpCAM+ cells and their lack of expression in EpCAM^-^ cells (**Fig. 4H**). Conversely, CAF-associated genes were enriched in EpCAM-, but not EpCAM+ cells (**Fig. 4H**). Together, these data provide transcriptional evidence that cancer cells and CAFs arise from distinct epithelial and mesenchymal lineages, respectively, in MCa-P1362 tumor explants.

**Figure 4.**
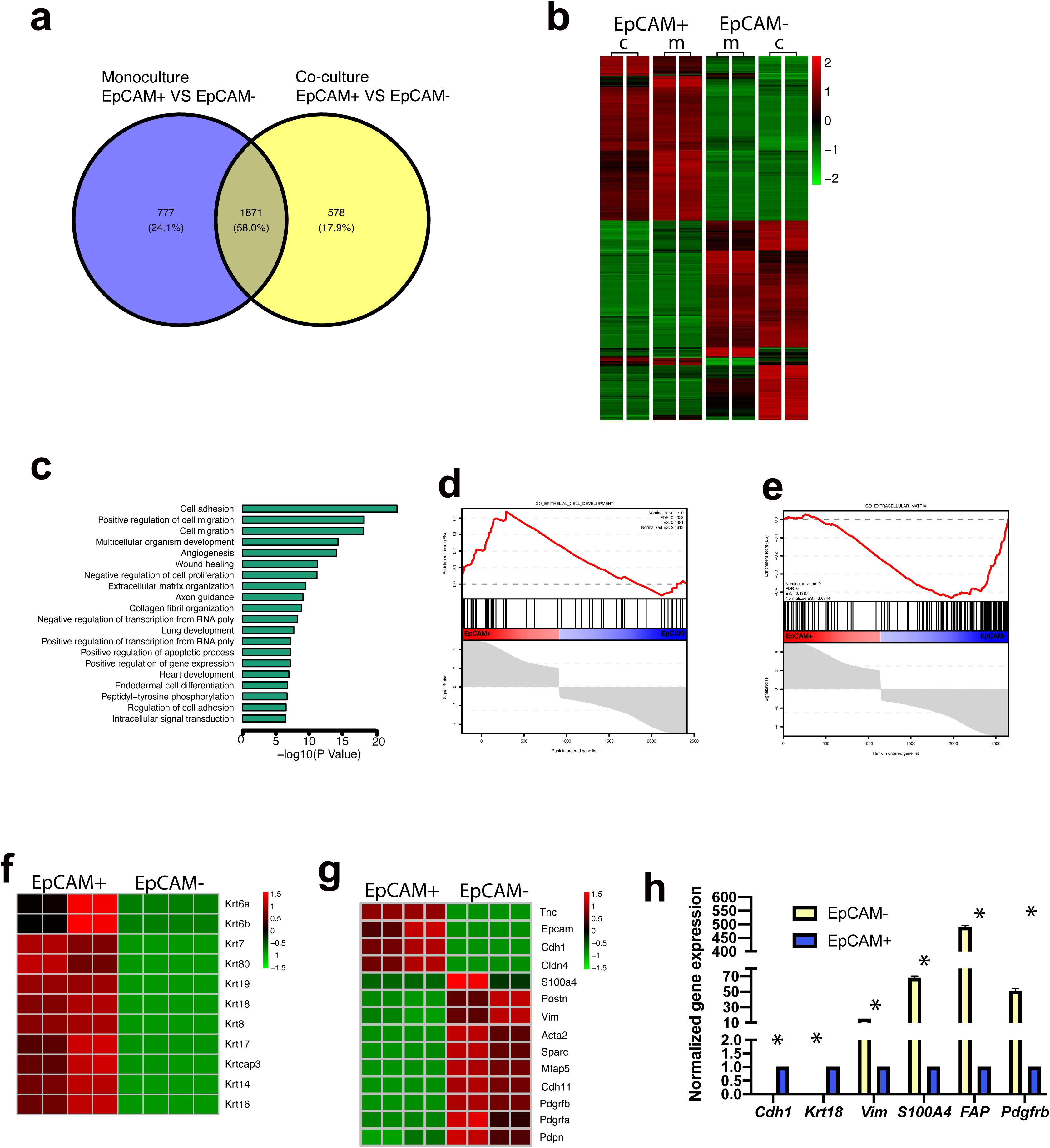
RNA sequencing of EpCAM+ and EpCAM-cells from MCa-P1362 tumors. (**A**) Venn diagram indicates number of differentially expressed genes (DEGs) between fluorescence-activity cell sorted (FACS) EpCAM- and EpCAM+ cells (grown in co-culture) and of DEGs between EpCAM+ and EpCAM-cells previously sorted and grown as separated monocultures. (**B**) Heatmap shows gene expression pattern of 3226 differentially expression genes between replicates of EpCAM- and EpCAM+ cells co-cultured (c) before collecting RNA of respective cell types or grown as monocultures (m) before harvest. (**C**) Gene ontology analysis of genes differentially expressed between EpCAM- and EpCAM+ cells. Gene set enrichment analysis results of epithelial cell development related genes (**D**) and extracellular matrix related genes (**E**) Differentially expressed keratin (**F**) and epithelial or mesenchymal-associated genes (**G**) between EpCAM+ and EpCAM-cells from bulk RNA-sequencing. (**H**) qPCR was used to measure relative expression levels of epithelial and mesenchymal genes in EpCAM+ and EpCAM-cells, respectively. Differential expression analysis was performed using the delta delta Ct method and genes of interest were normalized to β-actin.

### Enrichment of mesenchymal stem cells in MCa-P1362 primary culture

A comparison to the Single Cell Mouse Cell Atlas (scMCA) ^28^ revealed that the EpCAM^-^ cells showed a transcriptional profile similar to mesenchymal stem cells (MSCs) (**Fig. 5A**). We next used the ALDEFLUOR assay to detect aldehyde dehydrogenase (ALDH), a marker and regulator of cell stemness ^29^, in MCa-P1362 cells. Cells were incubated with ALDEFLUOR reagent alone or with both the ALDH inhibitor diethylaminobenzaldehyde (DEAB) and the ALDEFLUOR reagent. Using DEAB to establish the gating strategy, ALDEFLUOR staining showed that nearly 30% of the EpCAM-mesenchymal cells were stem cells (**Fig. 5B**).

**Figure 5.**
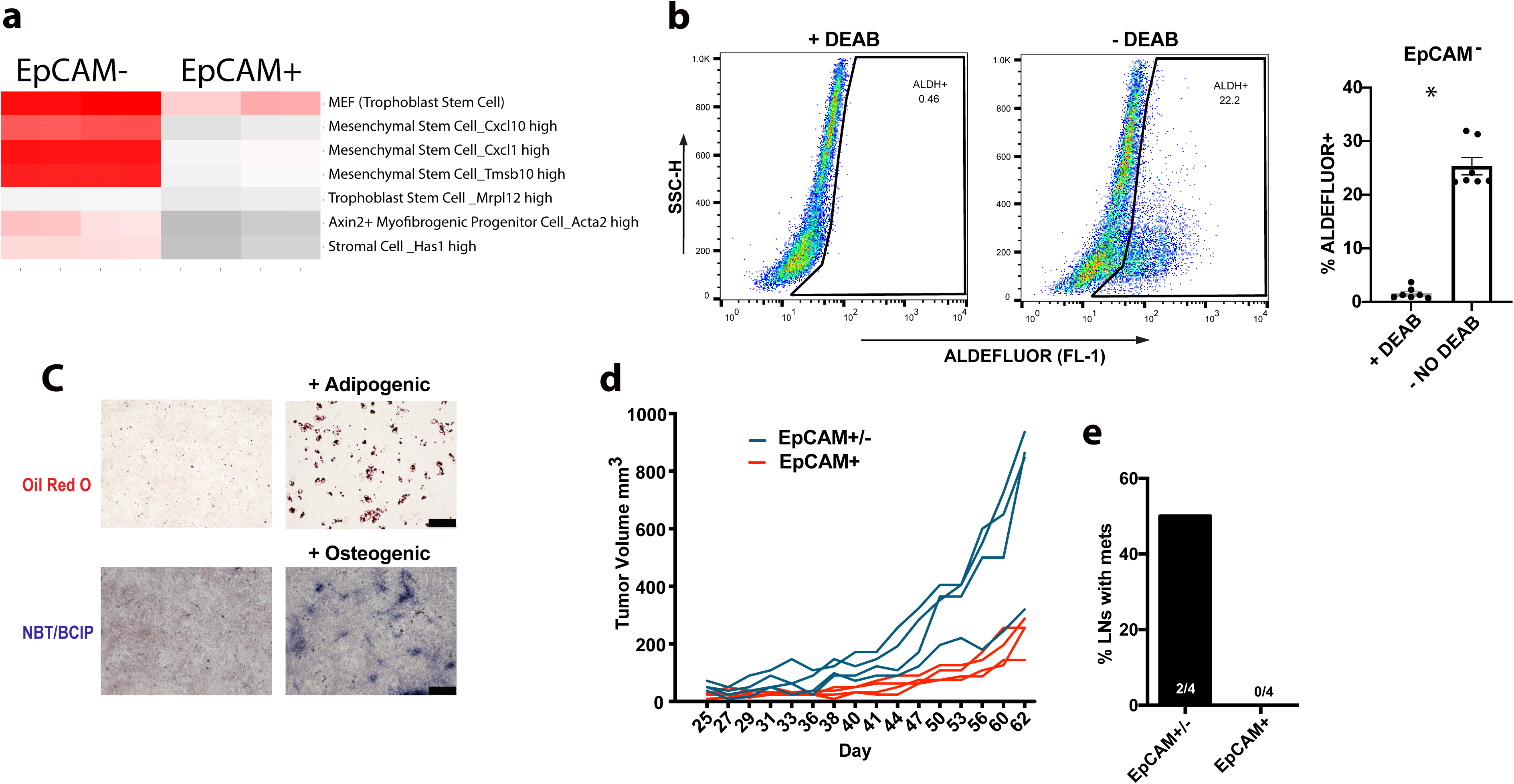
Characterization of mesenchymal cells from MCa-P1362 tumors. (**A**) Comparison of EpCAM- and EpCAM+ transcriptome to single-cell RNA sequencing transcriptomic atlas. (**B**) ALDEFLUOR-labeled EpCAM-cells were analyzed by flow cytometry with or without the ALDH inhibitor DEAB. A representative flow cytometry analysis of ALDEFLUOR fluorescence in EpCAM-cells is shown in the left panel. A gate was established based on the DEAB control and quantification of ALDEFLUOR+ cells is shown in the right panel. n=3 biological replicates * *p<* 0.0001. Significance was tested using two-tailed unpaired Student’s *t*-tests. (**C**) Adipogenesis of EpCAM-cells. Adipogenic media was used to cultivate EpCAM-cells and Oil Red O was used to stain for lipid droplets on after 7 days. Scale bars = 200 μm (**D**) Tumor growth curves in mice implanted with EpCAM+ cancer cells (red lines) or both EpCAM+ and EpCAM-mesenchymal cells (blue lines) from MCa-P1362 culture. Each line represents individual tumor growth. (**E**) Incidence of lymph node metastasis in mice implanted with EpCAM+ cells or both EpCAM+ and EpCAM-cells. n=4 in for D and E.

MSCs can differentiate into a variety of mesenchymal tissues including osteocytes, chondrocytes, and adipocytes ^30^. To measure the differentiation potential of EpCAM-cells from the MCa-P1362 line, EpCAM-cells were sorted and exposed to adipogenic or osteogenic differentiation medium to stimulate adipocyte and osteoblast formation, respectively. We found that EpCAM-cells differentiated into cells containing Oil Red O+ lipid droplets after incubation in adipogenic differentiation media (**Fig. 5C**) for 7 days. In addition, growth of EpCAM-cells in osteogenic media for 14 days led to the appearance of osteoblasts, as assessed by detection of alkaline phosphatase (**Fig. 5C**). The ability of MSCs to promote progression of both ER- and ER+ tumors ^31^ prompted us to investigate if EpCAM-cells could facilitate tumor growth. To this end, we implanted 1×10^6^ EpCAM+ cells or 1×10^6^ MCa-P1362 cells (EpCAM+ and EpCAM-cells) into recipient mice. Co-implantation of EpCAM+ and EpCAM-cells led to increased tumor growth compared to implantation with EpCAM+ cells alone (**Fig. 5D**). In addition, the incidence of lymph node metastasis was increased by co-implantation of EpCAM+ and EpCAM-cells (**Fig. 5E**). These data indicate that mesenchymal cells from MCa-P1362 culture promote cancer cell growth and metastasis.

### Enrichment of cancer stem cells in MCa-P1362 primary culture

EpCAM+ cells stained with ALDEFLUOR revealed a subset of ALDH+ cancer stem cells (CSCs) (**Fig. 6A**). Compared to MCF-7, EPCAM+ cells from MCa-P1362 primary culture contained significantly more CSCs. However, MCa-P1362 contained fewer CSCs than 4T1 cells, which are enriched in CSCs ^32^ (**Fig. 6B**). By Western blot, we investigated the protein expression of pluripotency markers Sox2, Oct4, and Nanog in MCa-P1362 and 4T1 cells. In addition, Sox9 and ALDH, also recognized as markers of breast cancer stem cells ^33^ were probed (**Fig. 6C**). 4T1 and MCa-P1362 cells showed similar expression of Oct4 and ALDH, while Nanog and Sox2 expression was lower in MCa-P1362 cells. Compared with 4T1 cells, MCa-P1362 cells expressed more Sox9 protein. To investigate the tumor-initiating capacity of MCa-P1362 CSCs, EpCAM+ ALDEFLUOR+, EpCAM+ ALDE-FLUOR-, EpCAM-ALDEFLUOR+, or EpCAM-ALDEFLUOR-cells were flow-sorted (**Fig. 6D**) and 5,000 cells from each group were injected into respective hosts. EpCAM+ ALDEFLUOR+ cells, but not other populations, were able to induce tumors in recipient mice (2 of 4) after 4 months of monitoring (**Fig. 6D**). These data together indicate that MCa-P1362 culture contains CSCs that are sufficient for tumor formation, independent of EpCAM-cells.

**Figure 6.**
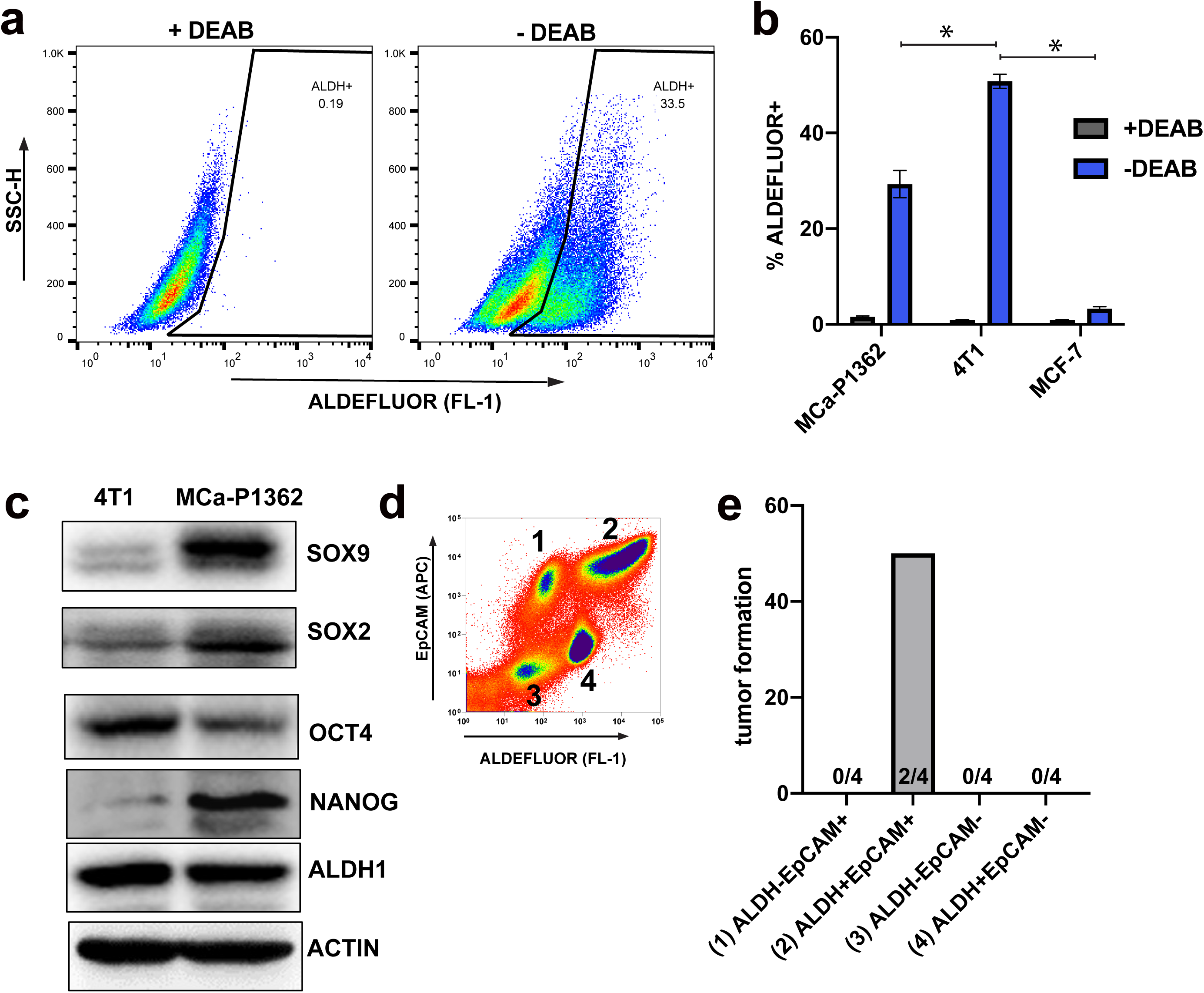
Characterization of cancer cells from MCa-P1362 tumors. (**A**) ALDEFLUOR-labeled EpCAM+ cells were analyzed by flow cytometry either with or without the ALDH inhibitor DEAB, and the left image shows a representative flow cytometry analysis of ALDEFLUOR fluorescence in EpCAM+ cells. Using the DEAB control, a gate was established based on the quantification of ALDEFLUOR+ cells in the bar graph. (**B**) Analysis of ALDEFLUOR-labeled cancer stem populations in EpCAM+ cells derived from MCa-P1362 culture as well as 4T1 and MCF-7 breast cancer cells. Cells incubated with DEAB were used as a control for the ALDEFLUOR label. n=3 biological replicates. Significance was tested using one-way ANOVA with Tukey’s honestly significant difference post hoc test. (**C**) Western blot showing SOX9, SOX2, OCT4, NANOG and ALDH proteins that are associated with cancer cell stemness. ß-Actin was used as a loading control. (**D**) Flow cytometry dot plot depicting the flow-sorting strategy for MCa-P1362 cells based on EpCAM and ALDEFLUOR labeling. (**E**) Bar graph depicting the incidence of tumor formation in animals injected with 5,000 cells from each sorted population. n=4/group.

## Discussion

Newer class of medicines such as PI3K inhibitors and CDK4/6 inhibitors have proven to be effective in treating advanced estrogen-receptor positive breast cancers when combined with endocrine therapies ^34^. Yet, resistance is developed to these current therapies. In the absence of ER+ mouse breast cancer models, it is difficult to identify additional targets and molecules for treating metastatic ER+ breast cancer. In addition, using mice to model human ER+ breast cancer has some limitations, such as the requirement of supplemental estrogen due to low levels of endogenous estrogen in mice and poor maintenance of the microenvironment of breast tumor growth. Further, studies of human tumor progression and treatment response in immunocompromised mice make it impossible to examine the role of the immune system in these experimental models.

As part of our current study, we characterized a novel hormone receptor-expressing tumor, P1362, derived from a spontaneous adenocarcinoma in an aged syngeneic Balb/c female breeder. Based on the expression of ERα, PR, HER2, and the proliferative index, P1362 tumors can be classified as a luminal B subtype, a common subtype diagnosed in patients and one that has poor prognosis ^35^. From P1362 transplanted tumors, we established an in vitro and *in vivo* MCa-P1362 tumor line. The MCa-P1362 tumor line shows 100% transplantability, consistent growth pattern *in vivo*, and about 50% metastasis rate to host lymph nodes, lungs, and livers. The MCa-P1362 cell line consisted of EpCAM+ cancer cells and EpCAM-mesenchymal cells that could be passaged indefinitely *in vitro*. The epithelial to mesenchymal transition has been shown to confer stemness to epithelial cells with the acquisition of mesenchymal-like characteristics ^36^. According to a previous study ^37^, mesenchymal and epithelial CSCs may constitute distinct, yet interconvertible subpopulations within breast tumors. While we cannot exclude this possibility, EpCAM-cells grow at a slower rate than EpCAM+ cells and EpCAM+ cancer cells eventually dominate the culture. We found that the EpCAM-, ALDEFLUOR+ population consists of mesenchymal stem cells, and we speculate that the EpCAM-, ALDEFLUOR-population consists of CAFs. Collectively, EpCAM-mesenchymal cells promoted tumor development and metastasis, potentially by providing cytokines and growth factors that directly act on the EpCAM+ cancer cells. It is also possible that EpCAM-cells may indirectly promote tumor growth by stimulating angiogenesis or by modulating anti-tumor immunity in the host. Finally, in this model, EpCAM-cells were also able to confer resistance to chemotherapy and endocrine therapy-induced cell death.

Serial passages of MCa-P1362 tumors *in vivo* resulted in faster tumor growth in this study and in others ^19^. Moreover, cultured MCa-P1362 cells reproducibly formed tumors when implanted into Balb/c mice but grew more slowly than tumors formed from serially passaged cancer cells. Importantly, MCa-P1362 tumors develop and progress in the absence of exogenous estrogen and with reduced endogenous hormone levels from ovariectomy. MCa-P1362 tumors. Moreover, we found that MCa-P1362 tumor growth rate is enhanced by the presence of EpCAM-cells. Dominance of EpCAM+ cells in MCa-P1362 cultures led to slower tumor development and progression compared to lower-passage cultures that contained more EpCAM-cells. Notably, MCa-P1362 tumor progression and metastasis are also significantly accelerated by exogenous estrogen, indicating that ERα in these tumors retains functional activity to induce tumor growth *in vivo*.

A limitation of this cell line is that it may not adequately capture the heterogeneity among breast cancer patients. Moreover, CAFs are heterogeneous, and each subset likely performs a specific function. The heterogeneity of EpCAM-cells was not investigated in this study. In conclusion, MCa-P1362 cancer cells implanted into syngeneic recipients overcome the limitations of immune deficiency and stroma mismatch. The established MCa-P1362 breast cancer model provides a platform for mechanistic studies of both cancer cell intrinsic and tumor microenvironment-mediated resistance to breast cancer therapies.

## Materials and Methods

### Cell lines and animals

The MCa-P1362 cell line was established from a spontaneous mammary tumor P1362 arose in a female syngeneic Balb/c breeder in the Cox-7 animal facility ^19–21^. MCF-7, BT-474, and 4T1 cells were purchased from ATCC. All cells were grown in high glucose DMEM medium containing 10% fetal bovine serum and were maintained in a 5% CO_2_-humidified incubator at 37°C.

### Tumor induction

0.5-1×10^6^ composite MCa-P1362 cells or EpCAM+ cells from MCa-P1362 culture in 50 µL volume of Hanks Balanced Saline Solution were implanted into the second, third, or fourth mammary fat pad (MFP) of 6–10-week-old female BALB/c mice. The animal experiment protocol was reviewed and approved by the Institutional Animal Care and Use Committee (IACUC) of Boston University and the Massachusetts General Hospital.

### Estrogen pellet implantation

Pellets containing 0.36 mg of 17-β estradiol (cat # SE-121) or placebo (cat # SC-111) were purchased from Innovative Research of America. Once mice were anesthetized, a small incision was made on the lateral side of the neck between the ear and the shoulder of the mouse. Surgical forceps were used to place pellets subcutaneously into the pocket and the wound was closed using Vetbond (3M, cat #1469SB).

### Ovariectomy

Bilateral ovariectomy was performed on 6-week-old mice under ketamine-xylazine anesthesia. A single incision was made near the ovary, allowing for careful extraction of the ovary through the muscle opening. Silk sutures were placed below the ovary to prevent bleeding and separation of the ovary from the uterus was achieved with a single cut. The muscle layer was closed using 5-0 polyester sutures (Ethicon) and the skin was closed with wound clips. Betadine was applied to the incision area to prevent infection. Subcutaneous administration of 0.1 mg/kg buprenorphine was provided to alleviate post-surgical discomfort. While mice were under anesthesia, 1×10^6^ MCa-P1362 cells were implanted into the 2^nd^ mammary fat pad. Tumor growth was assessed by caliper twice weekly.

### Western blot analysis

To extract the total protein, cells were harvested and lysed with cell lysis buffer (Cell Signaling Technology, cat #9803) with added inhibitor cocktail (Roche, cat # 11836153001). Protein concentration was determined using Bradford Assay Kit (Promega, USA) and equal amounts of proteins were denatured, separated by SDS-PAGE, and then transferred onto a PVDF membrane (Millipore, cat # IPVH00010). The membrane was then blocked with 5% nonfat milk for 1 hour at room temperature. Next, the membrane was incubated overnight with primary antibody at 4°C. Antibodies used for Western blot from Cell Signaling Technologies: SOX9 (cat # 82630), SOX2 (cat # 4900), Oct4 (cat # 2840), ALDHA1 (cat # 12035). Antibodies used for Western blot from Abcam: Nanog (cat # ab21624). Antibodies used for Western blot from Abclonal: Estrogen Receptor alpha (cat # A12976), Progesterone Receptor (cat # A0321), HER2/erbB2 (cat # A2071). Antibodies used for Western blot from Millipore Sigma: ß-actin (cat # A2228). Following primary antibody incubation, membranes were with HRP (horseradish peroxidase) conjugated secondary antibodies for 1 h at room temperature. Secondary antibodies were from Cell Signaling Technologies: Anti-mouse IgG, HRP-linked Antibody (cat # 7076), Anti-rabbit IgG, HRP-linked Antibody (cat # 7074). After that the blot signals were detected using an ECL Western blotting detection kit (Thermo Fisher, USA). Protein bands were visualized using iBright FL1500 Imaging System (Thermo Fisher, USA).

### Flow cytometry analysis of stem cells

The ALDH assay was performed in accordance with the manufacturer’s instructions for the ALDEFLUOR ^TM^ kit (Stem Cell Technologies, Cat # 01700). Cells were trypsinized, washed with PBS, and then stained with an APC-conjugated anti-EpCAM antibody (BioLegend, cat # 118214 (only mouse cells)) for 20 min at room temperature protected from light. After incubation, cells were washed with PBS. Next, cells were incubated with ALDEFLUOR reagent alone or with both the ALDH inhibitor diethylaminobenzaldehyde (DEAB) and the ALDEFLUOR reagent. After incubation for 45 min at 37°C according to manufacturer’s instructions, the cells were washed and analyzed by flow cytometry using a BD FACS Calibur (BD, USA). Further analyses of cellular populations were performed using FlowJo software version 10 (BD, USA).

### Immunofluorescence staining

After tissue harvesting and embedding with optimal cutting temperature compound, tissues were frozen at -80°C and 7-10-mm frozen sections were cut by a cryostat. Slides from fresh frozen samples were fixed in -20°C acetone for 10 minutes and then allowed to air-dry. Human tissue was processed as previously described ^38^. For cell staining, MCa-P1362 cells were grown in culture plates or in slide chambers (Electron Microscopy Services, cat # 70360-82), fixed with 4% PFA in PBS for 15 minutes at room temperature and then permeabilized with 0.1% triton-X buffer. To minimize non-specific antibody binding, tissue or cells were blocked in 5% normal donkey serum for 30-60 minutes at room temperature. Antibodies used for staining from Abclonal: Estrogen Receptor alpha (cat # A12976), Progesterone Receptor (cat # A0321), HER2/erbB2 (cat # A2071). Cancer cells were identified using a pan-cytokeratin antibody (Millipore Sigma cat # C2931). Mesenchymal cells were identified using an anti-mouse smooth muscle actin antibody (Invitrogen, cat # 53-9760-82) or an anti-human smooth muscle actin antibody (DAKO, cat # M0851). Anti-Ki67 antibody (Abcam cat # 15580) was used to identify proliferating cells. In the case of non-conjugated primary antibodies, secondary antibodies containing Alexa Fluor non-overlapping fluorophores (Jackson ImmunoResearch Laboratories) specific to the isotype of the primary antibody were added.

### Detection of metastases

To determine the incidence of lymph node metastasis, tumor-draining lymph nodes were pooled and serially sectioned until the frozen block was exhausted. After serial sectioning, the slides were stained with anti-cytokeratin (Millipore Sigma, cat # C2931) to detect metastases.

### Tamoxifen and estradiol treatment

4-OHT (Millipore Sigma, cat # H7904) and 17-β estradiol (Cayman Chemical Company, cat # 10006315) were dissolved in 100% ethanol and DMSO, respectively. For tamoxifen treatment, an equivalent number of cells in phenol free high glucose DMEM without phenol red (Gibco cat # 31-053-028) containing 10% charcoal-stripped FBS (Gibco cat # A3382101) were plated. After overnight cell attachment, tamoxifen or estrogen were diluted in the media at indicated concentrations.

For in vivo administration of tamoxifen (Millipore Sigma, cat # T5648), tamoxifen was dissolved in corn oil and 100 µg of tamoxifen was administered daily. Tumor sizes were measured every 2-3 days.

### Chemotherapeutic treatment

Paclitaxel and cisplatin were obtained from the Massachusetts General Hospital pharmacy and purchased from Fresenius Kabi, respectively, and diluted in cell culture media to final concentrations.

### RNA-Seq preparation

EpCAM+ and EpCAM-cells from MCaP-1362 culture were sorted at the Massachusetts General Hospital Flow Cytometry Core. Respective EpCAM- and EpCAM+ cells were either pelleted and immediately flash frozen after sorting or grown at 37°C/5% CO_2_ in a and passaged once before pelleting and flash freezing. Cell pellets were stored at -80°C until shipping to Admera Health (South Plainfield, NJ). RNA extraction and library construction was performed by Admera followed by 150 bp paired-end Illumina HiSeq sequencing.

### Bioinformatic analysis

FastQC ^39^ was used for the quality control of raw sequencing data. After quality control, Cutadapt ^40^ was used to remove the low-quality bases and adaptor contaminations. The quality of yield clean data was examined by FastQC software again. Next, Hisat2 ^41–43^ was used to align the clean data to mouse reference genome mm10 which we downloaded from Illumina iGenomes database. After data mapping, samtools ^44^ was used to manipulate SAM files and BAM files. HTSeq-count from HTSeq package ^45^ was used to count the number of reads that were aligned to the gene features. The differentially expressed genes were identified by edgeR ^46^ with cutoff of |log(fold change)| > 1 and p value < 0.01. GSEA analysis ^47^ was performed by GSEA software with standard genesets from MSigDB database. The visualization of GSEA results was performed by Rtoolbox from https://github.com/PeeperLab/Rtoolbox.

### Quantitative real-time Polymerase Chain Reaction (qPCR)

RNA was isolated with an RNeasy mini kit with QIAshredder to homogenize cells. The quality and quantity of RNA were determined spectrophotometrically at 260 nm and 280 nm using a NanoDrop ND-1000. Purified mRNA was reverse transcribed to cDNA using Quantitect Transcription kit (Qiagen, US, cat# 205313) according to the manufacturer’s instructions. Next, real-time PCR for was performed according to the manufacturer’s instructions using iTAQ Universal SYBR Green Supermix (BIO-RAD, cat# 172-5124) on a BIO-RAD CFX Connect™ Real-Time PCR Detection System to amplify the following genes:

Vimentin, (forward) 5’-CCCTGAACCTGAGAGAAACTAAC, (reverse) 5’-CTCTGGTCTCAACCGTCTTAATC ; E-Cadherin (forward) 5’-CTGCTGCTCC-TACTGTTTCTAC, (reverse) 5’-TCTTCTTCTCCACCTCCTTCT S100A4, (forward) 5’-ATTCAGCACTTCCTCTCTCTTG, (reverse) 5’-CACCCTCTTT-GCCTGAGTATT ; Fibroblast Activation Protein (forward) 5’-CTCCCTCGTCCAATTCAGTATC, (reverse) 5’-GTGGATCTCCTGGTCTTTGTT PDGFR-B, (forward) 5’-AAGGTGCTGGAGATGTTGAG, (reverse) 5’-GTTGTT-GCTGTCCGTGTTATG; Keratin 18 (forward) 5’-ACTCCGCAAGGTGGTAGATGA, (reverse) 5’-TCCACTTCCACAGTCAATCCA Beta-Actin (forward) 5’-CAGCCTTCCTTCTTGGGTATG, (reverse) 5’-GGCATA-GAGGTCTTTACGGATG Cell viability Cell Titer Glo viability assay Cells were plated in respective medium and were allowed to adhere overnight. The CellTiter-Glo Luminescent Cell Viability Assay (Promega, cat # G9241) was performed according to instructions. Briefly, half the medium was removed from the wells of the plates, and an equivalent volume of room temperature CellTiter-Glo Reagent was added. Plates were rocked for 2 min in the dark and incubated for an additional 8 min or longer (while protected from light) before reading luminescent output with a Varioskan LUX plate reader (Thermo Fisher, USA).

### MTT Assay

To each well, 10 µL of MTT (5 mg/mL) (Medchemexpress cat # 50-187-3393) was added, and the plates were incubated for another 3 hours. After removing the media from each well, DMSO was added to solubilize the formazan crystals. Absorbance at 570 nm was measured using a BioTek (USA) MicroQuant microplate reader.

### Differentiation of EpCAM-cells

EpCAM+ or EpCAM-cells were plated into 12 or 24-well plates. Upon 80% confluence, normal growth media was replaced with MesenCult™ (StemCell Technologies) adipo-genic (cat # 05507) or osteogenic (cat # 05504) differentiation media for every 3 days for 7-14 days for adipogenic differentiation; for osteoblast differentiation, differentiation media was changed every 3 days for 14-21 days. Cells were stained with Oil Red O (Sigma cat # O0625) and osteoblasts were detected with alkaline phosphatase chromogen; 5-bromo-4-chloro-3-indoxylphosphate/nitroblue tetrazolium chloride (BCIP/NBT) substrate (Abcam cat # ab7468).

### Cell purification using microbeads

A MACS-based isolation protocol was used to enrich EpCAM+ and EpCAM-cells from MCa-P1362 tumors. Briefly, cells were trypsinized to obtain a single cell suspension in 0.5% BSA in PBS. 10 µL CD326 MicroBeads (mouse; Miltenyi Biotec cat #130-105-958) were added (up to 1 × 10^7^ cells) and mixed with the cell suspension and incubated for 15 min at 4 °C protected from light. Cell suspensions were passed through an LS (Miltenyi Biotec cat #130-042-401) or MS (Miltenyi Biotec cat # 130-042-201) column attached to a MidiMACS Separator (Miltenyi Biotec cat # 130-042-302) on a MACS MultiStand (Miltenyi Biotec cat # 130-042-303). The cells collected in the flow-through and subsequent rinses were the unlabeled EpCAM-cells, while cells attached to the column were flushed from the column and collected as EpCAM+ cells.

### Statistical Analysis

Statistical analyses were performed using Prism 9 (GraphPad). Statistical significance was determined using two-tailed unpaired Student’s *t*-tests, one-way ANOVA with the Tukey’s honestly significant difference post hoc test or grouped two-way ANOVA (mixed effect).

## Acknowledgments

Fig. 1A, 1D were created with Biorender.com and exported with publication licenses.

## Declarations

### Ethical Approval

The Boston University and Massachusetts General Hospital Animal Care and Use Committees approved all animal studies.

### Authors’ contributions

P.H. and W.L. prepared figure 1. S.J. Z.W., and D.J prepared figures 2, 3, 5, and 6. P.L. prepared figure 4. D.J. wrote the main manuscript text. All authors reviewed the manuscript. D.J. and P.H. supervised the study.

### Availability of data and materials

For this study, the primary data supporting the results are presented in the paper and its Supplementary Information. Bulk RNA datasets generated for this study will be deposited to the Gene Expression Omnibus. Upon reasonable request, raw datasets are available for research purposes from the corresponding author.

