## Supplemental Figure and Table for "“Isolation and characterization of a novel hormone receptor positive mammary adenocarcinoma MCa-P1362 with stromal drivers of tumor growth, metastasis, and drug resistance”"

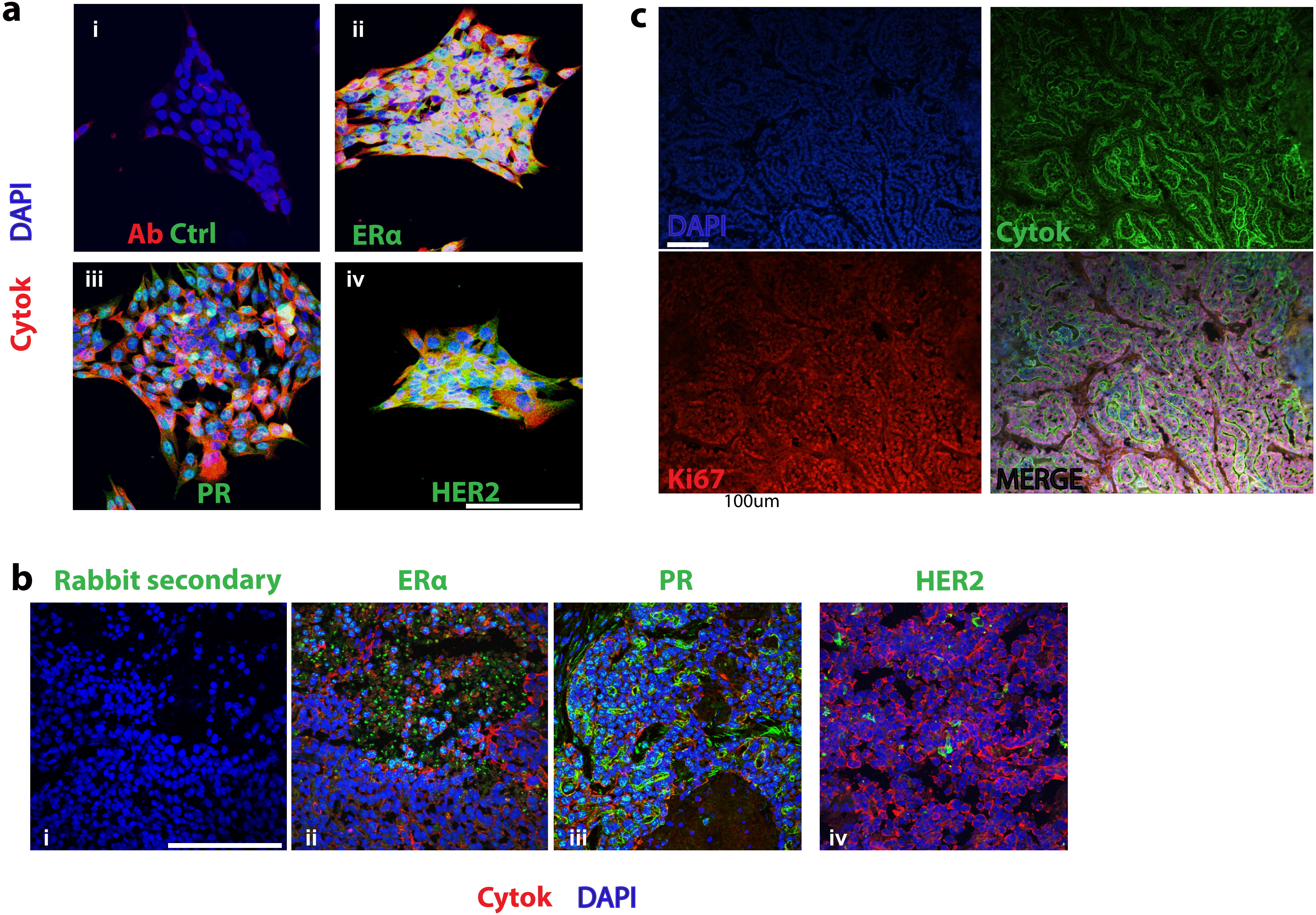
a
DAPI
Cytok
i
Ab Ctrl
ii
ERα
iii
PR
iv
HER2
c
DAPI
Cytok
Ki67
MERGE
100um
b
Rabbit secondary
ERα
PR
HER2
i
ii
iii
iv
Cytok DAPI

**Supplemental Figure 1**. Characterization of MCaP-1362 subtype. (**A**). MCaP-1362 cells were stained with IgG controls (i) or with primary antibodies against estrogen receptor alpha (ii), progesterone receptor (iii), or HER2 (iv). Cells were then stained with Alexa Fluor 488-conjugated secondary antibody (green). Anti-cytokeratin (red) was used to identify cancer cells. Cell nuclei are identified by DAPI (blue). Scale bar = 100 μm. (**B**)MCa-P1362 primary tumor tissue stained with Alexa Fluor 488-conjugated secondary antibody (i) and primary antibodies against estrogen receptor alpha (ii), progesterone receptor (iii), or HER2 (iv). Cancer cells were identified by anti-cytokeratin (red). Cell nuclei are identified by DAPI (blue). Scale bar = 100 μm (**C**) MCa-P1362 primary tumor tissue (day 49 post-implantation) stained with primary antibodies against Alexa Fluor 488-conjugated anti cytokeratin antibody and anti-Ki67. Tissue was then stained with Alexa Fluor 594-secondary antibody (red). Cell nuclei are identified by DAPI (blue). Scale bar = 100 μm.

-

| Tumor Line | Take rate* | Latent period (days)** | Growth time (days)*** |
|---|---|---|---|
| P1362 (F1-F5)<br>MCa-P1362 (C1-F4) | 95% (19 of 20)<br>100% (18 of 18) | 9.5 $\pm$ 7.7<br>8.4 $\pm$ 5.8 | 45.0 $\pm$ 18.4<br>39.6 $\pm$ 12.8 |

*Take rate: the number of mice in which the transplanted tumor tissue implanted and grew divided by the number of mice transplanted.

**Latent period: the number of days required for development of palpable MFP tumors post-transplantation.

***Growth time: the number of days required for the tumors to grow to 500 $\pm$ 50 $mm^3$.

**Table 1. Growth characteristics of P1362 & MCa-P1362 tumors in syngeneic Balb/c mice**
